# *Escherichia coli* growth under low pH, low temperature, and high osmotic stress conditions requires the *wzxE* flippase for the enterobacterial common antigen intermediate

**DOI:** 10.1101/2024.10.10.617665

**Authors:** Saki Yamaguchi, Kazuya Ishikawa, Kazuyuki Furuta, Chikara Kaito

## Abstract

Colanic acid and enterobacterial common antigen (ECA) are cell surface polysaccharides that are produced by many *E. coli* isolates. Colanic acid is induced under low pH, low temperature, and hyperosmotic conditions and is important in *E. coli* resistance to these stresses; however, the role of the ECA is unclear. In this study, we observed that knockout of the flippase *wzxE*, which converts ECA intermediates from the cytoplasmic side of the inner membrane to the periplasmic side, resulted in low pH sensitivity in *E. coli*. The *wzxE*-knockout mutant showed reduced growth potential and viable counts in the extracts of several vegetables (cherry tomatoes, carrots, celery, lettuce, and spinach), which are known to be low pH environments. A double knockout strain of *wzxE* and *wecF*, which encodes an enzyme that synthesizes an ECA intermediate, did not show sensitivity to low pH, nor did a double knockout mutant of *wzxE* and *wcaJ*, which encodes a colanic acid synthase. The *wzxE*-knockout mutant was sensitive to low temperature or hyperosmotic conditions, which induced colanic acid synthesis, and these sensitivities were abolished by the additional knockout of *wcaJ*. These results suggest that ECA intermediates cause *E. coli* susceptibility to low pH, low temperature, and high osmotic pressure in a colanic acid-dependent manner, and that *wzxE* suppresses this negative effect.

**Importance:** Polysaccharides covering bacterial cell surfaces, such as colanic acid, confer resistance to various stresses, such as low pH. However, the role of enterobacterial common antigens, carbohydrate antigens that are conserved throughout enterobacteria, in stress resistance is unclear. Our results suggest that lipid III enterobacterial common antigen, a substrate of flippase, causes sensitivity of *Escherichia coli* to low pH, low temperature, and high osmolarity in dependence on colanic acid synthesis, while *wzxE* inhibits this negative effect. The *wzxE*-knockout mutant was sensitive to crude vegetable extracts, suggesting that the creation of WzxE inhibitors could lead to new food poisoning prevention agents.

## Introduction

According to the WHO Foodborne Disease Burden Epidemiology Reference Group (2015), 35,000 people died from enteric diseases caused by enteropathogenic *Escherichia coli* in 2010, and *E. coli*-mediated foodborne diseases remain a problem today. Because *E. coli* is an enteric bacterium found in animal feces and urine, one of the most common causes of food poisoning is the contamination of crops by animal feces and manure in nature during the food processing stage. Food poisoning caused by pathogenic *E. coli* in cut vegetables and salads has become a problem. *Escherichia coli* is often exposed to low-pH stress in plant environments and many other stresses, such as low-temperature and drought stress. Therefore, it is important to understand the mechanisms underlying the resistance of *E. coli* to these stressors.

*Escherichia coli* resists stresses such as low pH by covering its cell surface with polysaccharides. Polysaccharides present on the surface of *E. coli* include O and K antigens, colanic acid, and enterobacterial common antigens (ECA). Because many strains commonly produce colanic acid and ECA, it is important to investigate the role of these molecules under stress. Colanic acid is secreted extracellularly as a high-molecular-weight polysaccharide and is known to be partly bound to the lipopolysaccharide (LPS) core. Colanic acid synthesis is enhanced by low pH (1–3), low temperature (4), osmotic stress (5), and desiccation (6), suggesting that colanic acid protects *E. coli* cells under various stressful conditions. In the EHEC strain O157:H7, mutations in the *wca* operon, which encodes colanic acid synthases, cause sensitivity to low pH (1).

ECAs are trisaccharide repeats; ECA_LPS_ bound to the LPS core and ECA_PG_ bound to phosphatidylglycerol exist on the cell surface, whereas ECA_CYC_ with a ring structure exists in the periplasm. Although ECA_CYC_ is involved in the outer membrane permeability barrier (7), other physiological roles of ECAs are poorly understood.

In this study, we examined the low pH sensitivity of strains deficient in genes involved in ECA synthesis to understand the physiological role of ECA. Our results revealed that *wzxE* knockout led to low pH sensitivity. WzxE is a lipid III ECA flippase that translocates lipid III ECA (an ECA synthesis intermediate) from the cytosol to the periplasmic side of the inner membrane (**Fig. 1A**). Accumulation of lipid III ECA, a substrate of WzxE, is lethal; however, in *E. coli wzxE*-knockout strains, lethal accumulation of lipid III ECA is prevented by the colanic acid intermediate flippase WzxC (**Fig. 1A**) and the O antigen intermediate WzxB (8–10). Although the *wzxE*-knockout strain has been reported to be susceptible to nalidixic acid and amikacin (11), its phenotype in low-pH environments has not been investigated. In this study, we aimed to elucidate the mechanism underlying the low pH sensitivity of the *wzxE*-knockout strain.

**Figure 1.**
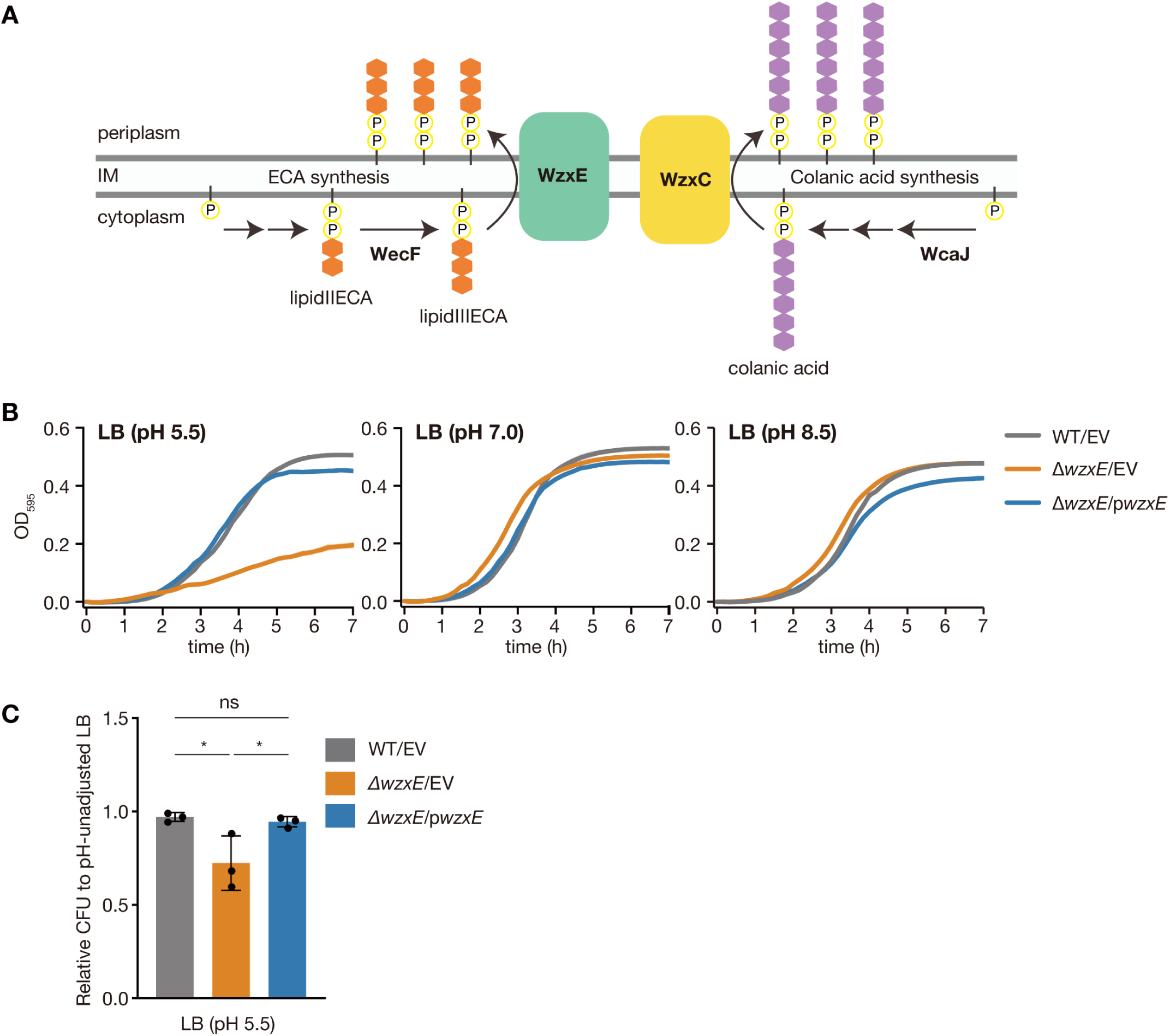
*wzxE*-knockout mutant is sensitive to low pH. A. Model diagram of the enterobacterial common antigen synthesis pathway and the colanic acid synthesis pathway. B. *Escherichia coli* wild-type strain transformed with pMW118 (WT/EV), *wzxE*-knockout mutant transformed with pMW118 (Δ*wzxE*/EV), and *wzxE*-knockout mutant transformed with pwzxE (Δ*wzxE*/pwzxE) were cultured in LB 50 mM MES (pH 5.5), LB 40 mM MOPS (pH 7.0), or LB 50 mM Tricine (pH 8.5), and the OD_595_ was measured for 7 h. C. Overnight bacterial cultures in LB medium were diluted and plated on LB agar medium or LB 50 mM MES (pH 5.5) agar medium, and the number of colonies formed was counted. Relative CFU on LB 50 mM MES (pH 5.5) agar against that on LB agar plate is shown. Data represent the mean ± SD of three independent experiments. One-way ANOVA with Tukey’s post hoc test p-values are provided (\**p* < 0.05).

## Results

### *wzxE-*knockout mutant is low pH sensitive

We first examined whether strains deficient in genes involved in ECA synthesis were sensitive to low pH and observed that the *wzxE*-knockout mutant showed reduced proliferative ability compared to the wild-type strain in LB medium adjusted to pH 5.5 (**Fig. 1B**). In contrast, the *wzxE*-knockout mutant showed no significant difference in growth in LB medium adjusted to pH 7.0 or pH 8.5 compared to the wild-type strain (**Fig. 1B**). Additionally, the *wzxE*-knockout mutant had fewer CFUs than the wild-type strain when an overnight culture grown in pH-unadjusted LB liquid medium was spread onto LB agar medium adjusted to pH 5.5 (**Fig. 1C**). The introduction of a plasmid carrying the *wzxE* gene into the *wzxE*-knockout mutant resulted in no significant decrease in proliferative ability or colony forming ability at pH 5.5 (**Fig. 1B, 1C**). These results suggest that *wzxE* is required for growth and colony formation under low pH conditions.

### *wzxE*-knockout mutant shows pH-dependent V8 sensitivity

In nature, plant environments are low-pH environments (12). We next investigated whether the *wzxE*-knockout mutant showed reduced proliferation in the V8 medium prepared from V8 vegetable juice, which is slightly acidic at pH 5.64. In the V8 medium, the *wzxE*-knockout mutant showed reduced proliferative ability compared to the wild-type strain (**Fig. 2A**). Additionally, the colony-forming ability of the *wzxE*-knockout mutant on the V8 agar medium was significantly reduced compared to that of the wild-type strain (**Fig. 2B**). The introduction of a plasmid carrying the *wzxE* gene into the *wzxE*-knockout mutant resulted in no significant reduction in growth capacity or colony-forming ability in V8 medium or V8 agar medium (**Fig. 2A, 2B**). These results suggest that the *wzxE*-knockout mutant is sensitive to the V8 medium environment.

**Figure 2.**
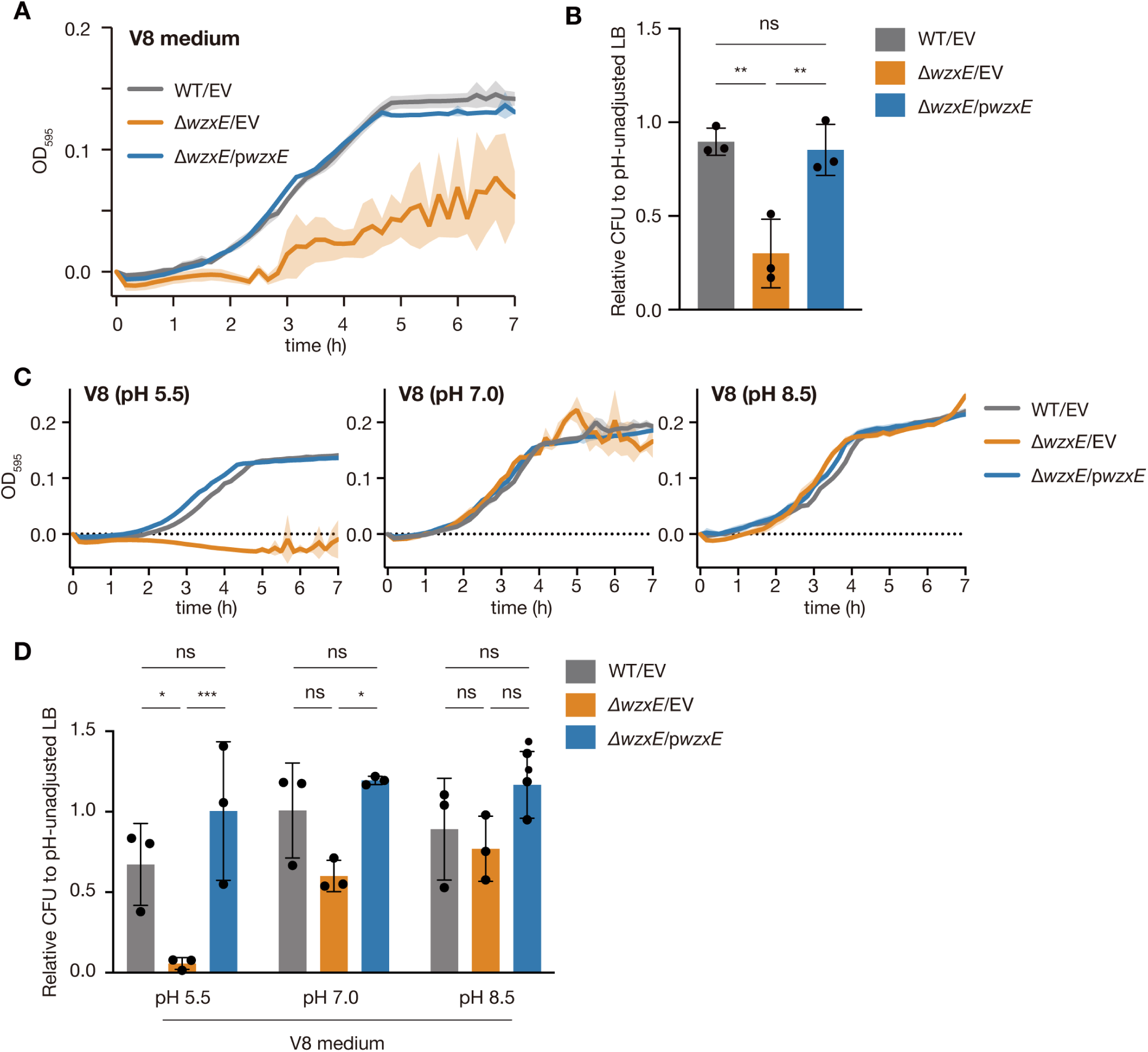
*wzxE*-knockout mutant is V8-sensitive. A. *Escherichia coli* wild-type strain transformed with pMW118 (WT/EV), *wzxE*-knockout mutant transformed with pMW118 (Δ*wzxE*/EV), and *wzxE*-knockout mutant transformed with pwzxE (Δ*wzxE*/pwzxE) were grown in V8 medium and the OD_595_ was measured for 7 h. Data represent the mean ± SD of three independent experiments. B. Overnight bacterial cultures in LB medium were diluted and plated on LB agar plate or V8 agar plate, and the number of colonies formed was counted. Relative CFU on V8 agar plate against that on LB agar plate is shown. Data represent the mean ± SD of three independent experiments. One-way ANOVA with Tukey’s post hoc test p-values are provided (** < 0.01). C. *E. coli* wild-type strain transformed with pMW118 (WT/EV), *wzxE*-knockout mutant transformed with pMW118 (Δ*wzxE*/EV), and *wzxE*-knockout mutant transformed with pwzxE (Δ*wzxE*/pwzxE) were cultured in V8 50 mM MES (pH 5.5), V8 40 mM MOPS (pH 7.0), or V8 50 mM Tricine (pH 8.5), and the OD_595_ was measured for 7 h. Data represent the mean ± SD of three independent experiments. D. Overnight bacterial cultures in LB medium were diluted and plated on LB agar medium, V8 50 mM MES (pH 5.5) agar medium, V8 40 mM MOPS (pH 7.0) agar medium, or V8 50 mM Tricine (pH 8.5) agar medium, and the number of colonies formed were counted. Relative CFU against that on LB agar plate is shown. Data represent the mean ± SD of three independent experiments. One-way ANOVA with Tukey’s post hoc test p-values are provided (\**p* < 0.05, *** < 0.001).

Next, we examined whether the V8 sensitivity of the *wzxE*-knockout mutant was pH-dependent. Increasing the pH of the V8 medium to 7.0 prevented a decrease in the growth potential of the *wzxE*- knockout mutant (**Fig. 2C**), while increasing the pH of the V8 agar medium restored the colony-forming ability of the *wzxE*-knockout mutant (**Fig. 2D**). These results suggest that the V8 sensitivity of the *wzxE*-knockout mutant is pH-dependent.

### *wzxE*-knockout mutant is sensitive to crude vegetable extracts

Given that the *wzxE*-knockout mutant was susceptible to the V8 medium, we sought to determine whether the *wzxE*-knockout mutant was susceptible to actual vegetables. To this end, we measured the number of viable bacteria after incubation in crude extracts of cherry tomato, carrot, celery, lettuce, and spinach. The results showed that the *wzxE*-knockout mutant had significantly lower viable counts than the wild-type strain in all vegetables (**Fig. 3**). In contrast, the *wzxE*-knockout mutant showed no significant difference in viable counts in saline compared to the wild-type strain (**Fig. 3**). The introduction of a plasmid carrying the *wzxE* gene into the *wzxE*-knockout mutant did not significantly reduce the viable counts in vegetables (**Fig. 3**). The pH of the vegetable crude extracts used in this study was 4.37 for cherry tomatoes, 6.45 for carrots, 6.23 for celery, 6.43 for lettuce, and 6.87 for spinach. A significant reduction in viable bacterial counts was observed in cherry tomatoes, which had a particularly low pH. These results suggest that the *wzxE*-knockout mutant is susceptible to these crude vegetable extracts.

**Figure 3.**
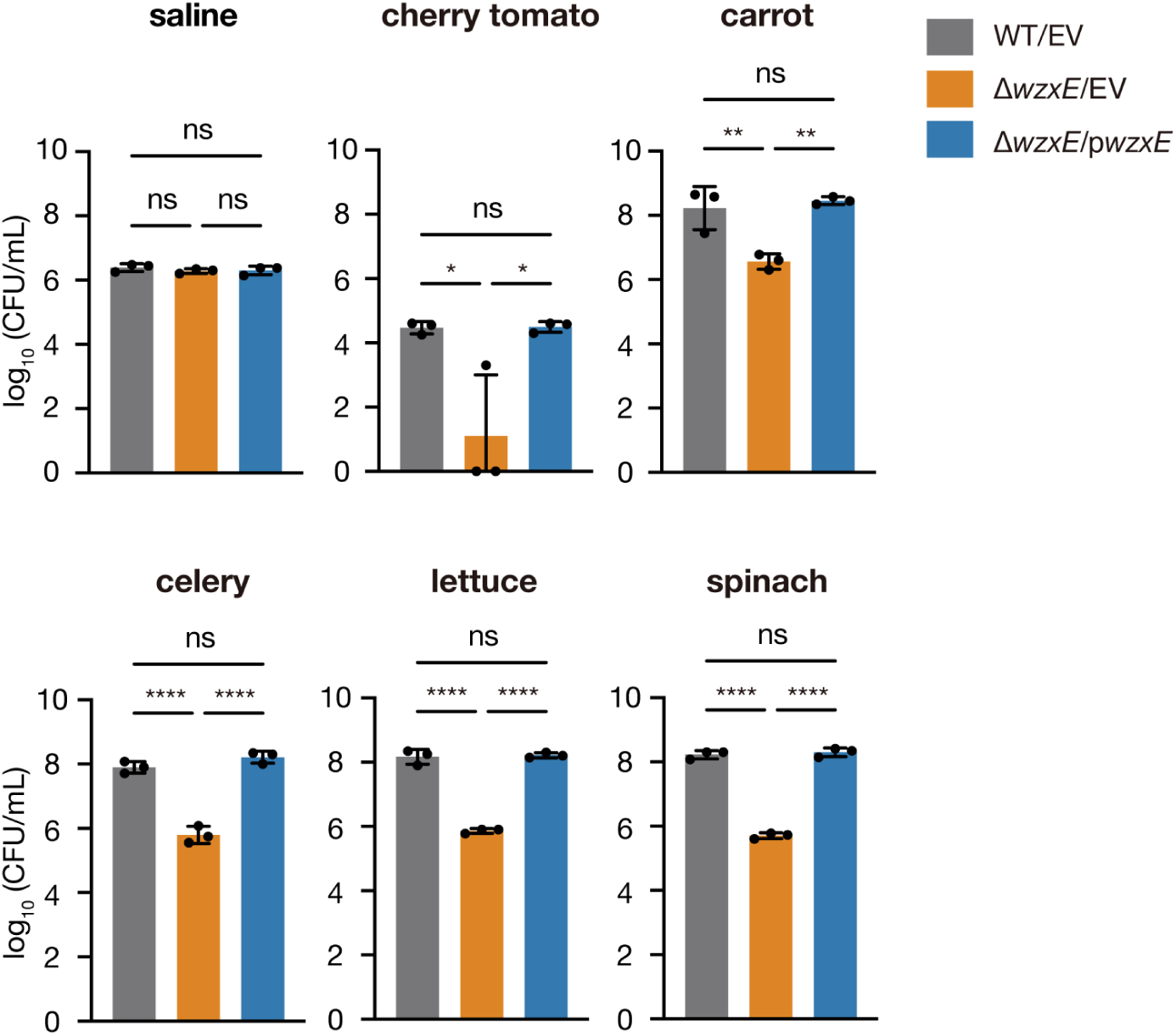
*wzxE*-knockout mutant is sensitive to vegetable crude extracts. Overnight bacterial cultures of the *E. coli* wild-type strain transformed with pMW118 (WT/EV), *wzxE*-knockout mutant transformed with pMW118 (Δ*wzxE*/EV), and *wzxE*-knockout mutant transformed with pwzxE (Δ*wzxE*/pwzxE) were inoculated into saline or crude extracts of small tomato, carrot, celery, lettuce, and spinach, and the viable counts were determined after 24 h of incubation at 25°C. The number of viable cells is shown on the vertical axis. Data represent the mean ± SD of three independent experiments. One-way ANOVA with Tukey’s post-hoc test p-values are provided (\**p* < 0.05, ** < 0.01, and **** < 0.0001).

### Lipid III ECA is involved in the low pH sensitivity of the *wzxE*-knockout mutant

The accumulation of lipid III ECA, a substrate of WzxE, on the cytoplasmic side of the inner membrane induces death in *E. coli* cells (8, 9). Therefore, we hypothesized that lipid III ECA is involved in the low pH sensitivity of the *wzxE*-knockout mutant. Consequently, we next investigated whether the pH sensitivity of the *wzxE*-knockout strain was abolished by knockout of *wecF*, a lipid III ECA synthase (**Fig. 1A**). The *wzxE*-knockout mutant showed reduced proliferative potential in low pH LB or V8, but the *wzxE*/*wecF* double-knockout mutant showed higher proliferative potential than the *wzxE* knockout mutant under these conditions (**Fig. 4A**). Furthermore, the *wzxE*/*wecF* double knockout mutant did not show reduced colony-forming ability on V8 agar medium compared to the wild-type strain (**Fig. 4B**). These results suggest that the lipid III ECA is involved in the low pH sensitivity of the *wzxE*-knockout mutant.

**Figure 4.**
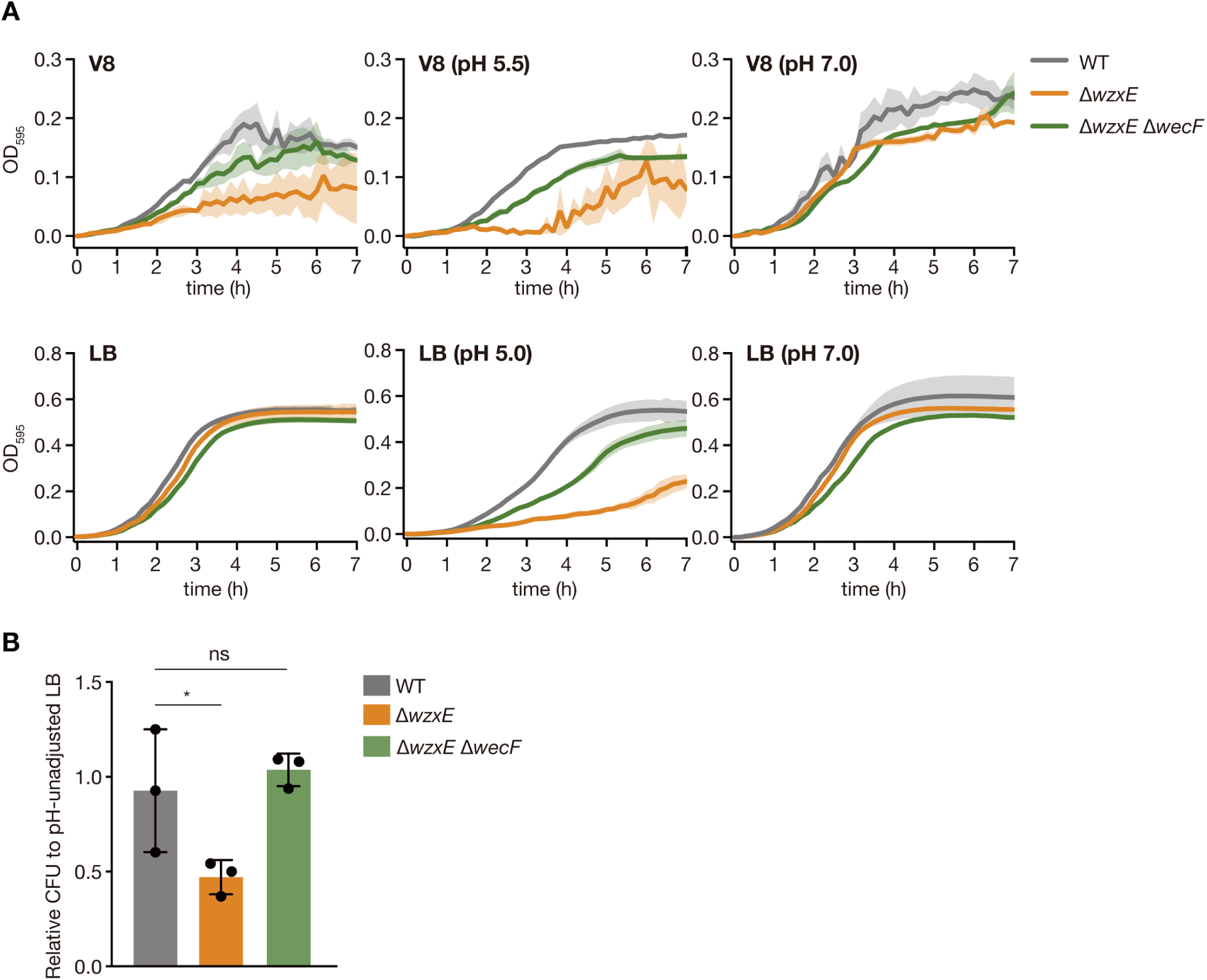
*wzxE*/*wecF* double-knockout mutant does not show low pH sensitivity. A. *Escherichia coli* wild-type strain (WT), *wzxE*-knockout mutant (Δ*wzxE*), and *wzxE*/*wecF* double-knockout mutant (Δ*wzxE* Δ*wecF*) were cultured in LB, LB 50 mM MES (pH 5.0), LB 40 mM MOPS (pH 7.0), V8, V8 50 mM MES (pH 5.5), or V8 40 mM MOPS (pH 7. 0), and the OD_595_ was measured for 7 h. Data represent the mean ± SD of three independent experiments. B. Overnight bacterial cultures in LB medium were diluted and plated on LB agar medium or V8 agar medium, and the number of colonies formed was counted. Relative CFU on V8 agar plate against that on LB agar plate is shown. Data represent the mean ± SD of three independent experiments. One-way ANOVA with Dunnett’s post hoc test p-values are provided (\**p* < 0.05).

### *wzxE*-knockout mutant contains numerous dead bacteria during growth under a low pH condition

To investigate the mechanism by which *wzxE* knockout reduced bacterial growth at low pH, the bacteria were treated with phosphate-buffered saline (PBS; pH 5.5) for 2 h, and the fluorescence of propidium iodide (PI) was measured using flow cytometry to determine the proportion of cell death. The results revealed no significant difference in the number of viable bacteria between the wild-type and *wzxE*-knockout strains treated with PBS (pH 5.5) (**Fig. 5A**). In contrast, when bacteria were cultured for 2 h in LB (pH 5.5), representing a growth-enabling condition, the number of dead bacteria was significantly higher in the *wzxE*-knockout mutant than in the wild-type strain (**Fig. 5B**). Additionally, the number of viable bacteria per turbidity was lower in the *wzxE*-knockout mutant than in the wild-type strain in LB medium (pH 5.5) (**Fig. 5C**). The reduction in viable bacteria per turbidity in the *wzxE*-knockout mutant was abolished by the additional knockout of *wecF* (**Fig. 5C**). These results suggest that *wzxE* knockout increases the proportion of dead bacteria in a lipid III ECA-dependent manner under low pH conditions that allow bacterial growth.

**Figure 5.**
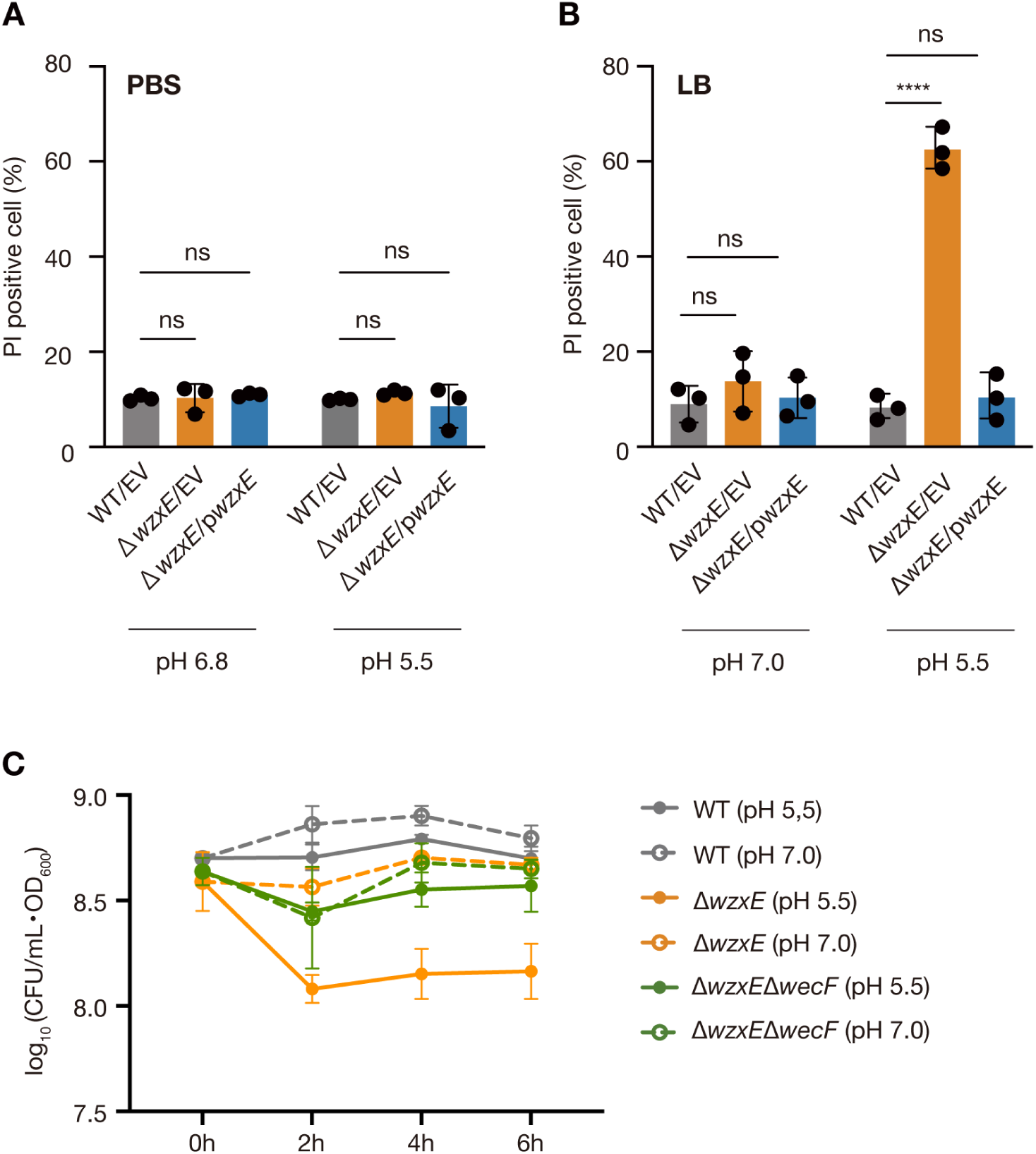
*wzxE*-knockout mutant shows an increase in dead bacteria in a low pH environment that allows bacterial growth. A. *Escherichia coli* wild-type strain transformed with pMW118 (WT/EV), *wzxE*-knockout mutant transformed with pMW118 (Δ*wzxE*/EV), and *wzxE*-knockout mutant transformed with pwzxE (Δ*wzxE*/pwzxE) were treated with PBS (pH 6.8) or PBS 50 mM MES (pH 5.5) for 2 h, followed by the addition of PI, and the fluorescence was measured using flow cytometry. Data represent the mean ± SD of three independent experiments. Two-way ANOVA with post hoc Dunnett’s test was performed (ns, *p* ≥ 0.05). B. Bacteria were aerobically cultured for 2 h in LB 40 mM MOPS (pH 7.0), or LB 50 mM MES (pH 5.5), stained with PI, and the fluorescence was measured using flow cytometry. Data represent the mean ± SD of three independent experiments. Two-way ANOVA with post hoc Dunnett’s test *p*-values are provided (\*\*\*\**p* < 0.0001). C. Overnight bacterial cultures were inoculated in LB 40 mM MOPS (pH 7.0) or LB 50 mM MES (pH 5.5) and the OD_600_ and viable counts were measured every 2 h. The vertical axis shows the CFUs per OD_600_. Data represent the mean ± SD of three independent experiments.

### Blockade of colanic acid synthesis abolishes the low pH sensitivity of the *wzxE*-knockout **mutant**

We hypothesized that *wzxE* knockout enhances ECA synthesis via *wecF* as a feedback response to ECA deficiency and worsens lipid III ECA accumulation associated with low pH sensitivity in the *wzxE*-knockout mutant. Our results revealed no significant differences in *wecF* expression between the wild-type strain and the *wzxE* knockout mutant at pH 5.5; however, we observed higher *wecF* expression in the *wzxE* knockout mutant than in the wild-type strain at pH 7 (**Fig. 6A**). Additionally, the *wecF* expression was not significantly different between pH 5.5 and 7 in the wzxE knockout mutant (**Fig. 6A**). These results show that *wecF* expression was not enhanced in the *wzxE*-knockout mutant under low pH conditions.

**Figure 6.**
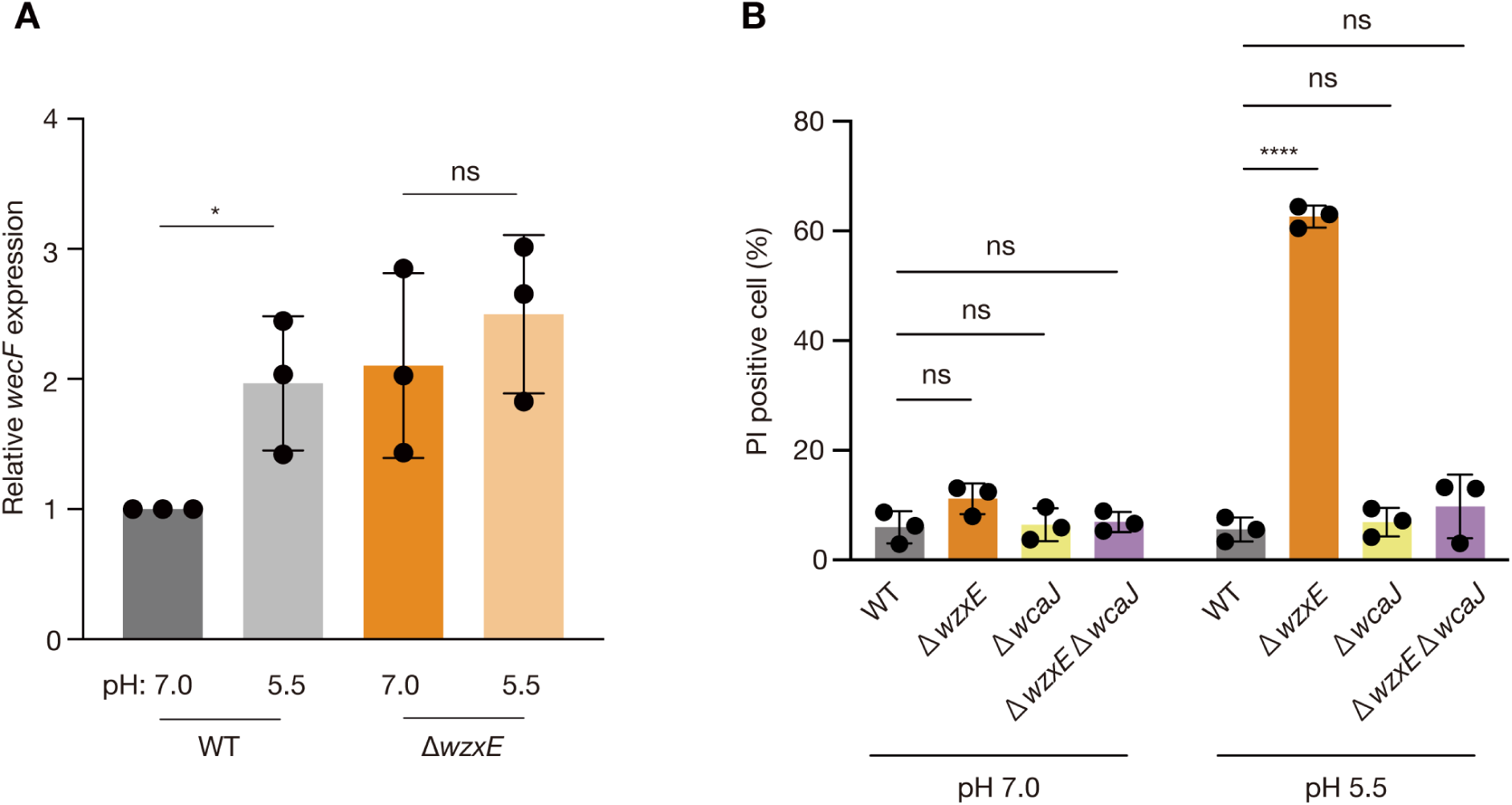
*wzxE*/*wcaJ* double knockout mutant does not show low pH sensitivity. A. Expression of *wecF* was determined using quantitative real-time PCR, with *gyrA* expression used as an internal standard. Data represent the mean ± SD of three independent experiments. Unpaired t test *p*-value is provided (\**p* < 0.05; ns, *p* ≥ 0.05). B. *Escherichia coli* wild-type strain (WT), *wzxE*-knockout mutant (Δ*wzxE*), *wcaJ-*knockout mutant (Δ*wcaJ*), and *wzxE*/*wcaJ* double knockout mutant (Δ*wzxE* Δ*wcaJ*) were aerobically cultured in LB 40 mM MOPS (pH 7.0) or LB 50 mM MES (pH 5.5) for 2 h, and the cells were stained with PI. The fluorescence was measured using flow cytometry. Data represent the mean ± SD of three independent experiments. Two-way ANOVA with post hoc Dunnett’s test p-values are provided (\*\*\*\**p* < 0.0001; ns, *p* ≥ 0.05).

The colanic acid intermediate flippase WzxC, and O antigen intermediate flippase WzxB are thought to flip lipid III ECA to prevent the lethal accumulation of lipid III ECA in *wzxE*-deficient strains (8–10). RcsA, a transcriptional activator of colanic acid synthesis, is upregulated in low-pH environments (2, 3). Based on these findings, we hypothesized that increased synthesis of colanic acid under low pH conditions causes WzxC to be occupied by colanic acid intermediates, reduces the amount of lipid III ECA flipped by WzxC, and increases the accumulation of lipid III ECA in the *wzxE*-knockout mutant, resulting in low pH sensitivity of the *wzxE*-knockout mutant. Experiments were performed using a *wcaJ*-knockout mutant, which lacks the first step of colanic acid synthesis, and a double-knockout mutants of *wzxE* and *wcaJ*. The *wcaJ*-knockout and double knockout mutants of *wzxE* and *wcaJ* showed no significant increase in dead bacteria after culture in LB (pH 5.5), whereas the *wzxE*-knockout mutant showed an increase in dead bacteria in LB (pH 5.5) (**Fig. 6B**). These results suggest that colanic acid synthesis is required for the low pH sensitivity caused by the *wzxE* knockout.

### *wzxE*-knockout mutant exhibits sensitivity to low temperature and high osmolality

Colanic acid is induced at low pH, low temperature, and high osmolality and promotes the survival of *E. coli* under these stresses (4, 5). We next examined whether the colanic acid synthesis-dependent susceptibility of the *wzxE*-knockout mutant was present not only at low pH, but also at low temperature and high osmotic stress. PI staining after 24 h incubation at 19°C, low-temperature stress, showed an increase in dead bacteria in the *wzxE*-knockout mutant compared to the wild-type strain (**Fig. 7A**). The double knockout mutants of *wzxE* and *wcaJ* showed no significant increase in dead bacteria after 24 h of incubation at 19°C (**Fig. 7A**). Additionally, PI staining after 2 h of incubation in LB (pH 7.0) adjusted with 300 mM NaCl, representing osmotic stress, showed an increase in dead bacteria in the *wzxE*-knockout mutant compared to that in the wild-type strain (**Fig. 7B**). The double knockout mutant of *wzxE* and *wcaJ* showed no significant increase in dead bacteria in the osmotic stress condition (**Fig. 7B**). In contrast, in LB adjusted with 100 mM NaCl, representing a low osmotic stress condition, the *wzxE*-knockout mutant showed a slight increase in dead bacteria compared to that in wild-type strain, but the difference was small compared to that in LB adjusted with 300 mM NaCl (**Fig. 7B**). These results suggest that the *wzxE*-knockout mutant is sensitive to low temperatures and osmotic stress in a colanic acid synthesis-dependent manner.

**Figure 7.**
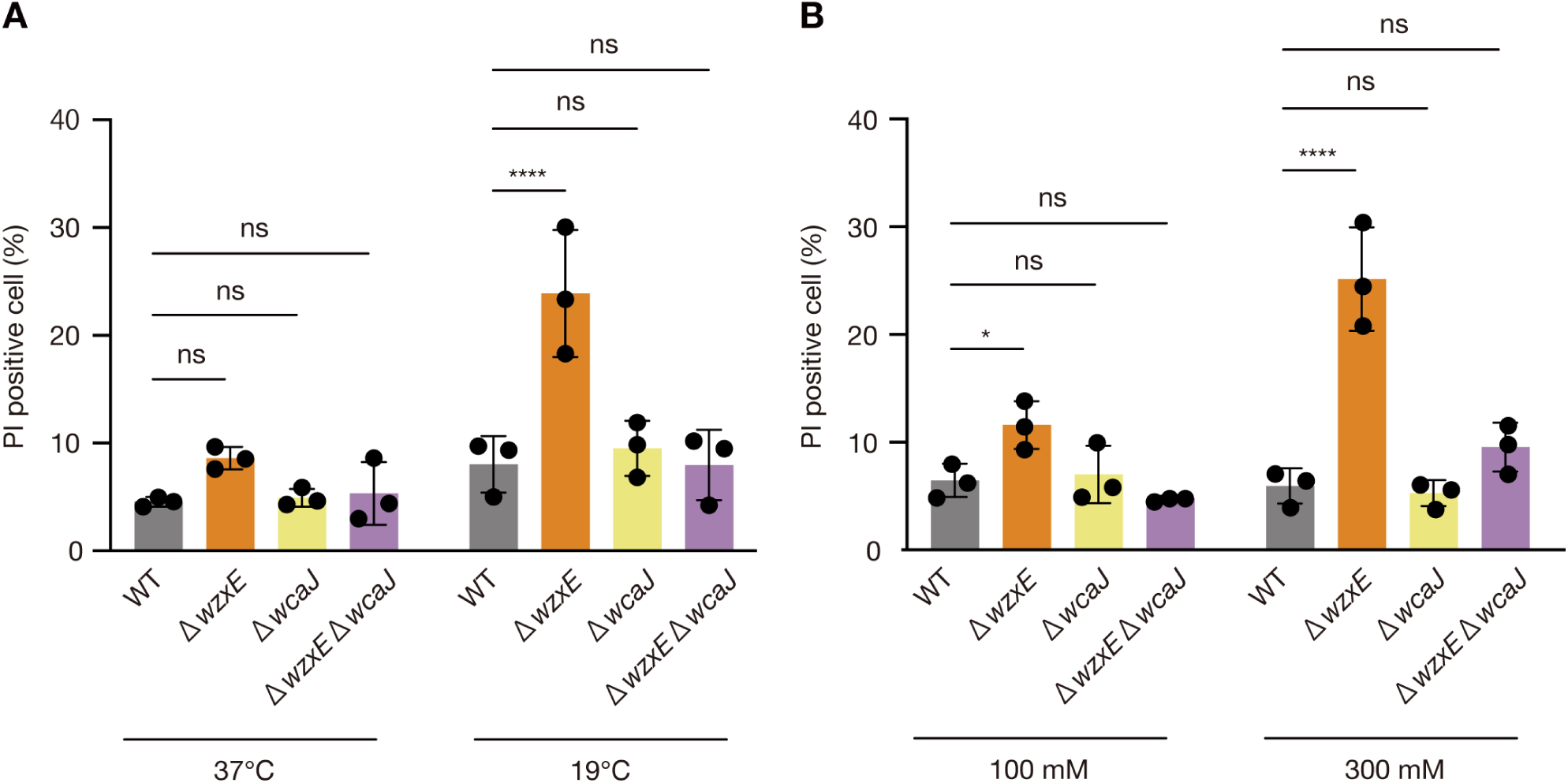
*wzxE*-knockout mutant is sensitive to low temperature and high osmolality. A. *E. coli* wild-type strain (WT), *wzxE*-knockout mutant (Δ*wzxE*), *wcaJ-*knockout mutant (Δ*wcaJ*), and *wzxE*/*wcaJ* double knockout mutant (Δ*wzxE* Δ*wcaJ*) were cultured in LB 40 mM MOPS (pH 7.0) for 24 h at 37°C or 19°C and then PI was added to stain cells. Two-way ANOVA with post hoc Dunnett’s test p-values are provided (\*\*\*\**p* < 0.0001; ns, *p* ≥ 0.05). B. *E. coli* wild-type strain (WT), *wzxE*-knockout mutant (Δ*wzxE*), *wcaJ-*knockout mutant (Δ*wcaJ*), and *wzxE*/*wcaJ* double knockout mutant (Δ*wzxE* Δ*wcaJ*) were aerobically cultured in LB 40 mM MOPS (pH 7.0) supplemented with 100 mM NaCl or 300 mM NaCl, and then PI was added and the fluorescence was measured using flow cytometry. Two-way ANOVA with post hoc Dunnett’s test *p*- values are provided (\**p* < 0.05; **** < 0.0001; ns, *p* ≥ 0.05).

## Discussion

In this study, we observed that the knockout of the lipid III ECA flippase *wzxE* caused *E. coli* susceptibility to low pH, low temperature, and osmotic stress. Furthermore, these stress susceptibilities depended on the lipid III ECA synthase *wecF* and colanic acid synthase *wcaJ*. This study is the first to demonstrate that the lipid III ECA flippase *wzxE* is required for resistance of *E. coli* to the stresses of low pH, low temperature, and osmotic pressure in a colanic acid synthesis-dependent manner.

Based on the results of this study, we hypothesized that the increased production of colanic acid in the *wzxE*-knockout mutant under low pH, low temperature, and high osmotic pressure conditions occupied WzxC and reduced the amount of lipid III ECA flipped by WzxC, resulting in the lethal accumulation of lipid III ECA (**Fig. 8**). To clarify this, future experiments are needed to quantify the amount of lipid III ECA and colanic acid in the *wzxE*-knockout mutant.

**Figure 8.**
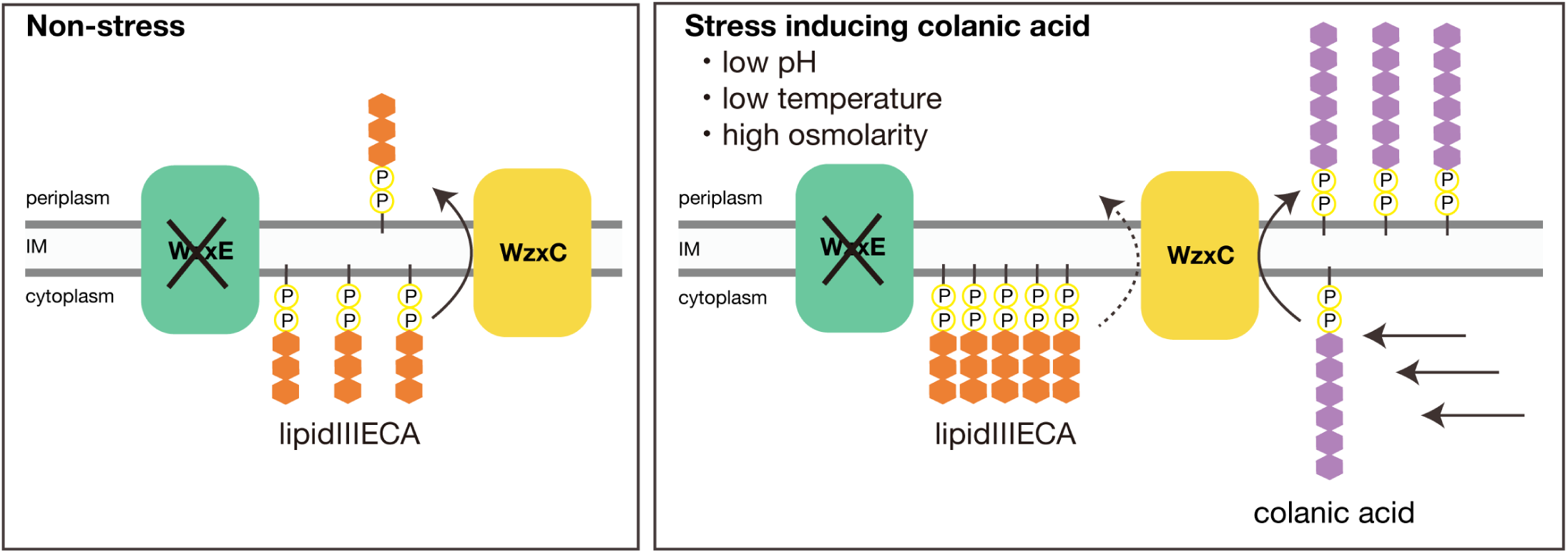
Mechanism underlying the sensitivity of the wzxE-knockout mutant against stresses that induce colanic acid synthesis. Under stress-free conditions that do not induce colanic acid synthesis, WzxC is assumed to prevent the accumulation of lipid III ECA in the *wzxE*-knockout mutant. However, under stress conditions, such as low pH, high osmotic pressure, and low temperature, which induce colanic acid synthesis, WzxC is occupied by colanic acid, lipid III ECA flipping by WzxC becomes less frequent, and lipid III ECA accumulates on the cytosolic side of the inner membrane, which may induce cell death.

The mechanism by which the accumulation of lipid III ECA on the cytoplasmic side of the inner membrane causes bacterial cell death is poorly understood. The non-lethal accumulation of ECA intermediates other than lipid III ECA suggests that lipid III ECA is particularly toxic. Mutants that accumulate ECA synthesis intermediates, such as the *wzxE*-knockout mutant, have been reported to have an abnormal cell shape, suggesting that the lipid carrier used for peptidoglycan synthesis, undecaprenyl phosphate, is trapped in the ECA synthesis pathway (13). Similar cell morphology abnormalities have been observed with the inhibition of O antigen and LPS core synthesis (14). The cell lysis of the *wzxE*-knockout mutant in a low-pH environment that allows bacterial growth may suggest a defect in cell division due to abnormal peptidoglycan synthesis.

The *E. coli* BW25113 strain used in this study is a K-12 strain that does not express the O antigen. Since the O antigen flippase *wzxB* can flip lipid III ECA in the *E. coli* K-12 strain (8–10), the phenotype of *wzxE* deficiency may change with or without O antigen expression and serotype. O-antigen carrying strains are of high public health importance because they are frequently isolated as causative agents of foodborne illnesses (15). Therefore, it is necessary to examine whether the mechanism described in this study applies to strains harboring the O antigen.

The decrease in the proliferative capacity and colony-forming ability of the *wzxE*-knockout mutant was more pronounced in the V8 medium (pH 5.5) than in the LB medium (pH 5.5) (**Figs. 1 and 2**). Additionally, there was no strict correlation between the pH of the crude vegetable extract and the number of viable bacteria after culturing in the crude vegetable extract (**Fig. 3**). Since the osmolality of the vegetable crude extracts used in this study was not significantly different from that of the saline solution, we speculate that factors other than pH and osmolality, such as the organic acids contained in vegetables, amplify the phenotype of the *wzxE*-knockout mutant in the vegetable environment.

Compared to the *wcaJ*-knockout mutant, which loses its ability to synthesize colanic acid, the *wzxE*-knockout mutant showed significantly higher sensitivity to low pH, low temperature, and hyperosmotic stress (**Figs. 6 and 7**). Furthermore, the *wzxE*-knockout mutant showed reduced proliferative capacity for all vegetable extracts analyzed in this study (**Fig. 3**). In addition, WzxE homologs are not found in mammals, including humans, but are widely distributed and highly conserved in Enterobacteriaceae (16). The family Enterobacteriaceae contains several bacterial species that cause food poisoning, including *Shigella*, *Salmonella*, and *Yersinia* species. These findings suggest that the development of inhibitors against WzxE leads to new food poisoning prevention agents that break the bacterial cycle between the plant environment and humans by inhibiting the growth of *E. coli* in low-pH environments such as the plant environment and intestinal tracts.

## Materials and Methods

### Bacterial strains and culture conditions

*Escherichia coli* BW25113 and its gene-knockout mutants were cultured on an LB agar medium, and their colonies were incubated aerobically at 37°C in an LB liquid medium. *Escherichia coli* strains transformed with pMW118 were cultured on LB agar medium containing 100 µg/mL. The details of the bacterial strains and plasmids used in this study are listed in Table 1.

**Table 1.** Bacterial strains and plasmids used in this study.

| Strain or plasmid | Genotype or characteristics | Reference |
| --- | --- | --- |
| Strains |  |  |
| BW25113 | *rrnB*, Δ*lacZ*4787, *HsdR*514<br>$\Delta$(*araBAD*)567, $\Delta$(*rhaBAD*)568, *rph*-1 | NBRC |
| JW3766 | BW25113 Δ*wzxE*; Km<sup>r</sup> | Baba *et al.*(22) |
| JW2032 | BW25113 Δ*wcaJ*; Km<sup>r</sup> | Baba *et al.*(22) |
| JM109 | Host strain for cloning | Takara Bio |
| Plasmids |  |  |
| pMW118 | Low-copy-number plasmid; Amp<sup>r</sup> | Nippon Gene |
| pMW118-*wzxE* | pMW118 with *wzxE*, Amp<sup>r</sup> | This study |
| pKD46 | A temperature sensitive plasmid expressing RED recombinase; Amp<sup>r</sup> | CGSC, Datsenko *et al.* (18) |
| pCP20 | A temperature sensitive plasmid expressing FLP recombinase; Amp<sup>r</sup> | CGSC Datsenko *et al.* (18) |

### Preparation of V8 medium

A slightly modified version of a previously published method (17) was used. 340 mL of V8 vegetable juice (Campbell) was stirred with 5 g of CaCO_3_. The mixture was then centrifuged at 2590 *g* for 10 min, and the supernatant was diluted 5-fold with Milli-Q water and autoclaved to obtain V8 medium. V8 agar medium was prepared by adding 20 g/mL agar to the unautoclaved V8 medium, followed by autoclaving. The pH-adjusted medium was prepared by adding buffer and adjusting the pH using NaOH.

### Genetic manipulation

The *wzxE* or *wcaJ* gene-knockout mutants were generated *via* transduction with P1 phage *vir* using JW3766 or JW2032 as phage donors and BW25113 as the recipient strain. The *wzxE* gene-knockout mutant was transformed with pCP20, resulting in a marker-less *wzxE-*knockout mutant. Transduction with P1 phage *vir* was performed using the marker-less *wzxE-*knockout mutant as a recipient strain and JW2032 as a phage donor to generate a *wzxE*/*wcaJ* double-knockout mutant. To construct the complementation plasmid, a DNA fragment containing *wzxE* was amplified using PCR with BW25113 genomic DNA as a template and oligonucleotide primer pairs (**Table 2**). The DNA amplification product was inserted at the XbaI and HindIII sites of pMW118 to obtain pMW118- wzxE. To generate the *wzxE*/*wecF* double-knockout mutant, the *wzxE* and *wecF* genes was deleted in BW25113 using the one-step inactivation method (18).

**Table 2.**
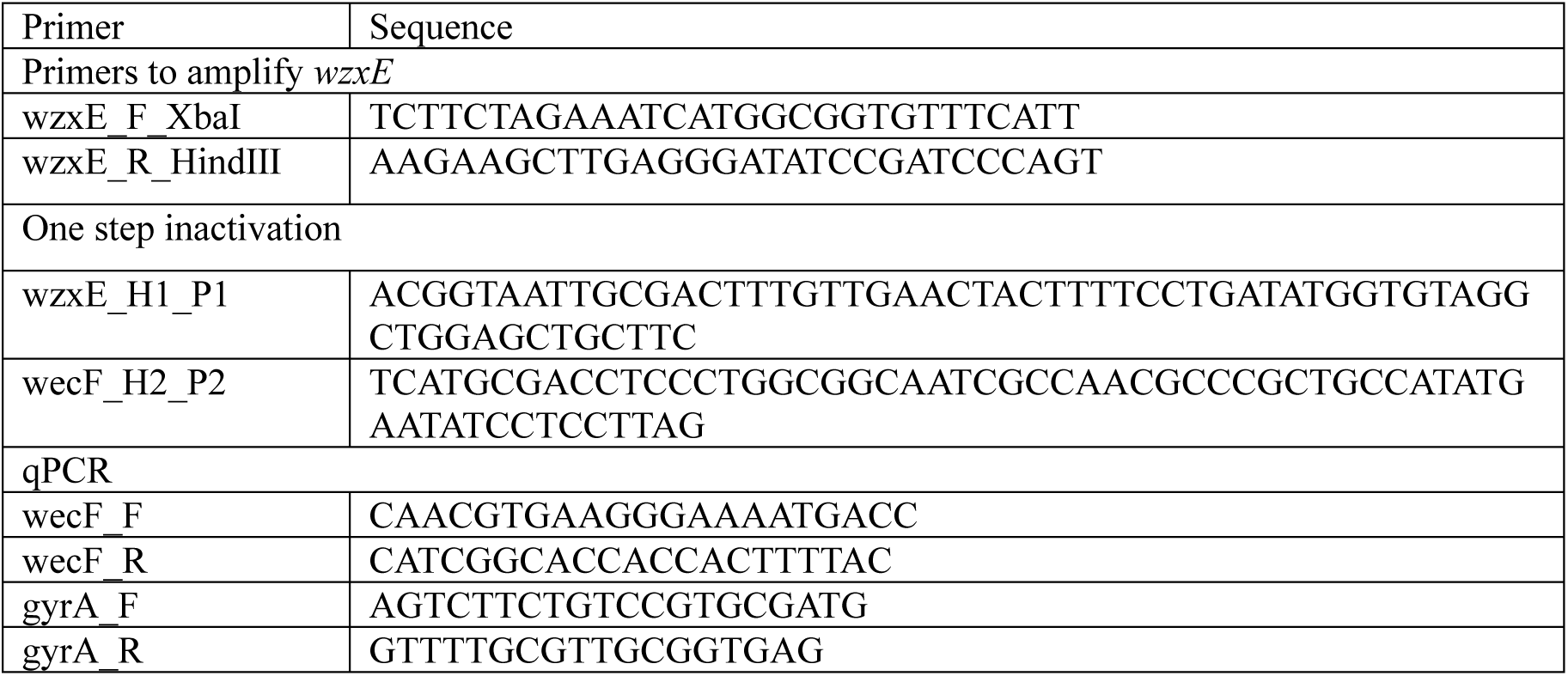
Primers used in this study.

### Growth curve

Briefly, 2 µL of an overnight culture of *E. coli* was added to 100 µL of the medium in a 96-well microplate and covered with a plastic seal. The OD_595_ was measured at 37°C for 7 h in a microplate reader.

### Colony forming ability

Dilutions of overnight cultures of *E. coli* were spread on an agar medium, and the number of colonies formed after incubation at 37°C was counted. The incubation times differed depending on the medium used: overnight for LB, LB 40 mM MOPS (pH 7.0), LB 50 mM tricine (pH 8.5), V8 40 mM MOPS (pH 7.0), and V8 50 mM tricine (pH 8.5); 2 nights for LB 50 mM MES (pH 5.5); and 3 nights for V8 50 mM MOPS (pH 5.5).

### Determination of the number of viable bacteria in the plant crude extracts

Crude vegetable extracts were prepared by crushing each vegetable and filtering it through gauze. Overnight bacterial cultures were added to 100 µL of vegetable crude extract supplemented with ampicillin and incubated at 25°C overnight. The culture medium was then serially diluted and applied to the LB agar medium, and CFU was measured after overnight incubation. The same procedure was performed for samples incubated overnight at 25°C in 0.9% NaCl as controls.

### Flow cytometry

A slightly modified version of a previously published method (19) was used for the analysis. Bacterial cells from overnight cultures were collected via centrifugation, suspended in PBS or PBS 50 mM MES (pH 5.5), the OD_600_ was adjusted, and incubated at 37°C for 2 h. Alternatively, overnight cultures were inoculated in LB 50 mM MES (pH 5.5) or LB 40 mM MOPS (pH 7.0), the OD_600_ was adjusted and incubated at 37°C for 2 h, and suspended in PBS. Subsequently, PI (7.5 µg/mL) was added, and the fluorescence was measured using flow cytometry after incubation at 37°C for 15 min in the dark. For cold stress, the bacteria were incubated in LB 40 mM MOPS (pH 7.0) at 19°C for 24 h, stained with PI, and analyzed using flow cytometry. For osmotic stress, bacteria were cultured for 2 h in LB 40 mM MOPS (pH 7.0) adjusted to NaCl 100 mM or 300 mM, PI stained, and then measured using flow cytometry.

### Quantitative PCR

The total RNA from *E. coli* was analyzed using a slightly modified version of a previously reported method (20, 21). *E. coli* cells were cultured in LB 50 mM MES (pH 5.5) or LB 40 mM MOPS (pH 7.0) for 2 h. Following incubation, 1.8 mL of the culture was mixed with 200 µL of ethanol containing 5% phenol by vortexing, centrifuged at 21500 × *g* for 2 min, and the precipitates were frozen in liquid nitrogen. The bacteria were dissolved in 200 µL of lysis buffer (TE buffer, 1% lysozyme, 1% SDS) and incubated at 65°C for 2 min. RNA was extracted using the RNeasy Mini Kit (QIAGEN, Tokyo, Japan), treated with DNase I, and used as a template for cDNA synthesis using SuperScript II reverse transcriptase and a random hexamer. Quantitative PCR was performed using the synthesized cDNA as a template using oligonucleotide primer pairs (**Table 2**) and KOD SYBR pPCR Mix (TOYOBO, Osaka, Japan). QuantStudio 3 (Thermo Fisher Scientific, Tokyo, Japan) was used for the assay, with *gyrA* used as the internal control.

### Statistical analysis

All statistical analyses were performed using GraphPad PRISM software.

## Acknowledgement

We gratefully thank Division of Instrumental Analysis, Department of Instrumental Analysis & Cryogenics, Advanced Science Research Center, Okayama University for technical assistance. This study was supported by Japan Society for the Promotion of Science (JSPS) Grants-in-Aid for Scientific Research (Grants 22K14892, 23K24131, 23K06130, 24K01760, 24K21872). This study was also supported by the Takeda Science Foundation (to C.K.), the Ichiro Kanehara Foundation (to C.K.), and the Ryobi Teien Memory Foundation (to K.I. and C.K.). We thank the National BioResource Project-E. coli (National Institute of Genetics, Japan) for providing the *E. coli* Keio collection.

## References

1. Mao Y, Doyle MP, Chen J. 2001. Insertion mutagenesis of wca reduces acid and heat tolerance of enterohemorrhagic Escherichia coli O157:H7. J Bacteriol 183:3811–5.

2. Kannan G, Wilks JC, Fitzgerald DM, Jones BD, Bondurant SS, Slonczewski JL. 2008. Rapid acid treatment of Escherichia coli: transcriptomic response and recovery. BMC Microbiol 8:37.

3. Maurer LM, Yohannes E, Bondurant SS, Radmacher M, Slonczewski JL. 2005. pH regulates genes for flagellar motility, catabolism, and oxidative stress in Escherichia coli K-12. J Bacteriol 187:304–19.

4. Navasa N, Rodríguez-Aparicio L, Ferrero M, Monteagudo-Mera A, Martínez-Blanco H. 2013. Polysialic and colanic acids metabolism in Escherichia coli K92 is regulated by RcsA and RcsB. Biosci Rep 33.

5. Sledjeski DD, Gottesman S. 1996. Osmotic shock induction of capsule synthesis in Escherichia coli K-12. J Bacteriol 178:1204–6.

6. Ophir T, Gutnick DL. 1994. A role for exopolysaccharides in the protection of microorganisms from desiccation. Appl Environ Microbiol 60:740–5.

7. Mitchell AM, Srikumar T, Silhavy TJ. 2018. Cyclic Enterobacterial Common Antigen Maintains the Outer Membrane Permeability Barrier of Escherichia coli in a Manner Controlled by YhdP. mBio 9.

8. Rick PD, Barr K, Sankaran K, Kajimura J, Rush JS, Waechter CJ. 2003. Evidence that the wzxE gene of Escherichia coli K-12 encodes a protein involved in the transbilayer movement of a trisaccharide-lipid intermediate in the assembly of enterobacterial common antigen. J Biol Chem 278:16534–42.

9. Kajimura J, Rahman A, Rick PD. 2005. Assembly of cyclic enterobacterial common antigen in Escherichia coli K-12. J Bacteriol 187:6917–27.

10. Marolda CL, Tatar LD, Alaimo C, Aebi M, Valvano MA. 2006. Interplay of the Wzx translocase and the corresponding polymerase and chain length regulator proteins in the translocation and periplasmic assembly of lipopolysaccharide o antigen. J Bacteriol 188:5124–35.

11. Girgis HS, Hottes AK, Tavazoie S. 2009. Genetic architecture of intrinsic antibiotic susceptibility. PLoS One 4:e5629.

12. Bridges MA, Mattice MR. 1939. Over two thousand estimations of the ph of representative foods. Digestive Diseases and Sciences 6:440–449.

13. Jorgenson MA, Kannan S, Laubacher ME, Young KD. 2016. Dead-end intermediates in the enterobacterial common antigen pathway induce morphological defects in Escherichia coli by competing for undecaprenyl phosphate. Mol Microbiol 100:1–14.

14. Jorgenson MA, Young KD. 2016. Interrupting Biosynthesis of O Antigen or the Lipopolysaccharide Core Produces Morphological Defects in Escherichia coli by Sequestering Undecaprenyl Phosphate. J Bacteriol 198:3070–3079.

15. Nataro JP, Kaper JB. 1998. Diarrheagenic Escherichia coli. Clin Microbiol Rev 11:142–201.

16. Böttger EC, Jürs M, Barrett T, Wachsmuth K, Metzger S, Bitter-Suermann D. 1987. Qualitative and quantitative determination of enterobacterial common antigen (ECA) with monoclonal antibodies: expression of ECA by two Actinobacillus species. J Clin Microbiol 25:377–82.

17. Anonymous. 1974. Mycological Society of America. In Stevens RB (ed), Mycology guidebook, University of Washington Press.

18. Datsenko KA, Wanner BL. 2000. One-step inactivation of chromosomal genes in Escherichia coli K-12 using PCR products. Proc Natl Acad Sci U S A 97:6640–5.

19. Yamamoto R, Ishikawa K, Miyoshi Y, Furuta K, Miyoshi SI, Kaito C. 2024. Overexpression of diglucosyldiacylglycerol synthase leads to daptomycin resistance in Bacillus subtilis. J Bacteriol doi:10.1128/jb.00307-24:e0030724.

20. Sanchez-Vazquez P, Dewey CN, Kitten N, Ross W, Gourse RL. 2019. Genome-wide effects on Escherichia coli transcription from ppGpp binding to its two sites on RNA polymerase. Proc Natl Acad Sci U S A 116:8310–8319.

21. Shirakawa R, Ishikawa K, Furuta K, Kaito C. 2023. Knockout of ribosomal protein RpmJ leads to zinc resistance in Escherichia coli. PLoS One 18:e0277162.

22. Baba T, Ara T, Hasegawa M, Takai Y, Okumura Y, Baba M, Datsenko KA, Tomita M, Wanner BL, Mori H. 2006. Construction of Escherichia coli K-12 in-frame, single-gene knockout mutants: the Keio collection. Mol Syst Biol 2:2006.0008.

